# Mitochondrial DNA Imputation Accuracy and Its Application in a Southern African Mitochondrial Genome-wide Association Study

**DOI:** 10.64898/2025.12.11.693816

**Authors:** Dayna Croock, Caitlin Uren, Marlo Möller, Desiree C. Petersen

## Abstract

Southern African populations harbor exceptional mitochondrial genetic diversity and include the earliest diverging mitochondrial haplogroups. Yet, they remain significantly underrepresented in genetic studies and reference databases. This underrepresentation raises concerns about the performance and generalizability of mitochondrial imputation tools in populations with complex demographic histories. In this study, we conducted a comprehensive evaluation of the publicly available MitoImpute reference panel to assess its accuracy and suitability for use in southern African cohorts. We examined imputation performance for mitochondrial DNA (mtDNA), the impact of imputation on haplogroup assignment, and the extent to which population-specific variants were recovered by the panel. As a proof-of-principle application, we used the MitoImpute reference panel and imputation workflow to merge five southern African tuberculosis (TB) case–control mtDNA datasets and perform a mitochondrial genome-wide association study (miWAS) of TB susceptibility. Although MitoImpute performed well in predicting mitochondrial haplogroups and accurately imputing a subset of mtDNA variants, substantial population-specific variation was not captured. No significant associations with TB susceptibility were identified, highlighting key limitations of array-based mtDNA genotyping and imputation in highly diverse populations. Together, these findings underscore the need for improved reference panels enriched for African mitochondrial diversity and provide practical guidelines for mitochondrial data harmonization and analysis in multi-ancestry cohorts.

## 1. Introduction

The mitochondrial genome serves as an important marker in population genetic studies of modern human origins, evolution, and migration patterns^1^. Given its unique characteristics of maternal inheritance, lack of recombination and high mutation rate, mitochondrial DNA (mtDNA) is a powerful tool for inferring maternal ancestry. Consequently, mtDNA has played a central role in elucidating the origins of the most recent maternal common ancestor which, based on current genetic evidence, is thought to have emerged in Africa approximately 200 000 years ago^2^. As anatomically modern humans migrated out of Africa and populated the rest of the world, new mitochondrial lineages arose. Closely-related maternal lineages with common single-nucleotide polymorphisms (SNPs), defined as haplogroups, display region-specific signatures and are useful in studying population structure^3^. Southern African populations exhibit some of the highest levels of genetic diversity worldwide^4^, a pattern that extends to the mitochondrial genome^5,6^. The earliest diverging lineages from the most recent maternal common ancestor, haplogroups L0d and L0k, are found exclusively in southern Africa and are present at the highest frequencies in Khoe-San populations^6^. These haplogroups are also observed in populations that historically received substantial maternal contributions from the Khoe-San, such as the South African Coloured (SAC) population^7^. Representing the earliest branches in the human mitochondrial phylogenetic tree, L0d and L0k possess the greatest genetic diversity of all global haplogroups^6,8^. Despite their well-documented genetic diversity, southern African populations remain underrepresented in mitochondrial sequence databases and genetic studies, consistent with the broader underrepresentation of African populations in genomic research^9,10^.

The primary function of the mitochondria is to produce energy through oxidative phosphorylation. Additionally, mitochondria play essential roles in metabolism^11^. Considering their role in several fundamental cellular processes, numerous studies have sought to identify mitochondrial mutations and single-nucleotide variants (mtSNVs) associated with complex diseases such as type 2 diabetes^12–14^, cardiovascular disease^15,16^ and other metabolic diseases^17^. Two large-scale consortia, the UK Biobank (UKBB)^18^ and the CHARGEmtDNA+ Consortium^17^, have provided workflows for quality control (QC), imputation, dataset harmonisation and the analysis or meta-analysis of mtDNA genotypes associated with common traits and identified numerous mtSNVs associated with complex diseases. However, mtSNV association studies performed in southern African populations are scarce. Given the exceptional genetic diversity observed in these populations, applying published workflows requires particular caution, as population structure in southern Africa is considerably more complex than in more homogeneous cohorts such as the UKBB, where mtDNA haplogroups are primarily European^18^. However, including southern African populations in mtDNA genetic studies could facilitate the discovery of novel mtSNVs associations with complex diseases, as well as uncover mtSNVs unique to African or southern African populations.

Imputation is a necessary step in cohort harmonisation, genome-wide association studies (GWASs) and meta-analyses^19^. Moreover, imputing untyped variants in the mitochondrial genome can yield haplogroup assignments that better approximate the true haplogroups^20^. Publicly available reference panels and imputation tools designed specifically for the mitochondrial genome are scarce. MitoIMP (released in 2019) is a tool for inferring missing nucleotides in low-coverage human mitochondrial genomes^21^. However, as it requires input sequences in single-sequence FASTA format, it is not easily adaptable for imputation in mtDNA datasets derived from genotype arrays. In 2021, the first publicly available mtDNA imputation reference panel was published, together with an accompanying imputation algorithm, MitoImpute^22^. Since the MitoImpute algorithm integrates the IMPUTE2 genotype imputation software^23^, it can be applied to array-based mtDNA datasets. The MitoImpute reference panel includes 36 960 complete mitochondrial genomes from diverse populations, representing all major haplogroups worldwide^22^. However, some haplogroups are disproportionately represented in the panel compared to others (Supplementary Figure 1). Specifically, haplogroup H, which is predominantly associated with European ancestry, is significantly overrepresented in the panel relative to other global haplogroups. The developers of MitoImpute evaluated the performance of their reference panel using *in silico* microarrays derived from the 1000 Genomes (1000G) Phase 3 Project^22,24^. However, the 1000G cohort notoriously lacks adequate representation from southern African haplogroups (Supplementary Figure 2). Given the exceptional genetic diversity observed in southern African mitochondrial genomes, the suitability of this panel for accurate imputation in this population must be rigorously assessed. In this study, we evaluate the performance of the MitoImpute reference panel in two southern African cohorts and apply it to a TB case–control genotype dataset for a mitochondrial GWAS (miWAS) of TB susceptibility, illustrating its utility for harmonising multi-platform data in a cohort with complex population structure. To our knowledge, this is the first comprehensive evaluation of a publicly available mtDNA imputation panel in a southern African population. Moreover, this study provides the first miWAS of TB susceptibility and, to our knowledge, the first mitochondrial genome-wide association study performed in any southern African cohort.

## 2. Materials and methods

### 2.1. Evaluation of imputation performance and reliability

Three cohorts were used to assess imputation performance and accuracy: (A) a set of 110 complete mtDNA sequences from individuals of the L0d haplogroup within a SAC population (unpublished data); (B) a set of 164 complete mtDNA sequences from a southern African cohort, representing macrohaplogroups L0 – L5^25^; and (C) a subset of 687 complete mtDNA sequences from the 1000G Phase 3 cohort belonging to macrohaplogroups L0 – L5. The distribution of haplogroups in each cohort is shown in Figure 1. Notably, the 1000G Phase 3 cohort does not contain any individuals belonging to the L0d haplogroup.

**Figure 1.**
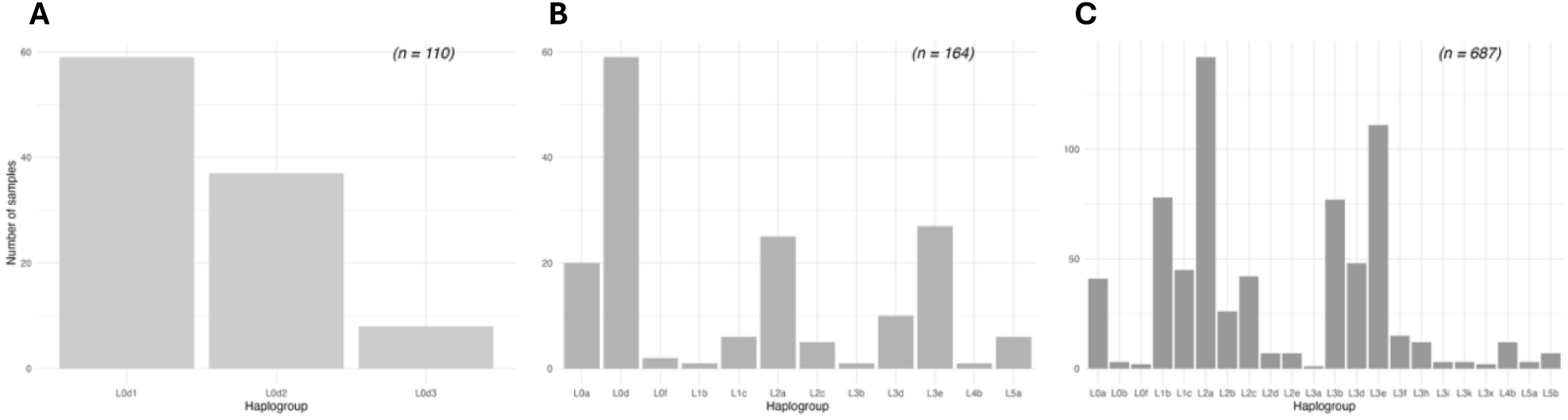
Distribution of haplogroups for **(A)** the southern African L0d cohort, **(B)** the southern African L-macrohaplogroup cohort and **(C)** the 1000 Genomes Phase 3 L-macrohaplogroup cohort. Haplogroups were assigned using complete mitochondrial sequences and Haplogrep v3. Cohort sample sizes are shown in parentheses on the graph backgrounds.

For this analysis, *in silico* genotype microarray data was generated for each cohort by extracting mtSNVs from the complete mtDNA sequences. For each cohort, we assessed imputation performance across eight commercially available arrays (Table 1), which have been assessed for their ability to assign African mitochondrial haplogroups accurately^26^. Genotyped mtSNV positions for each array were obtained from the Infinium manifest files for Illumina BeadChip arrays and the annotation files for the Thermo Fisher Scientific Affymetrix and Axiom arrays (Supplementary Data: Table 1).

**Table 1.**
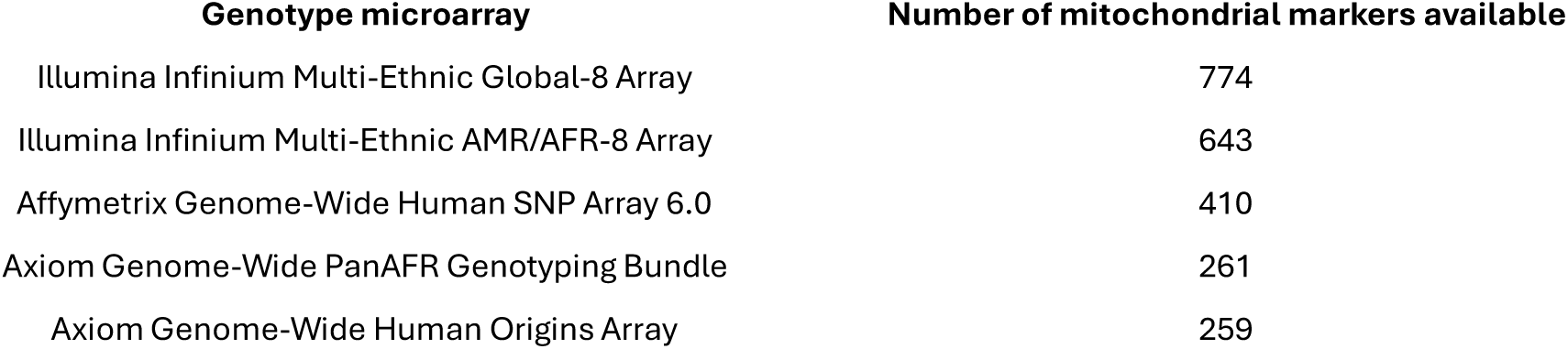
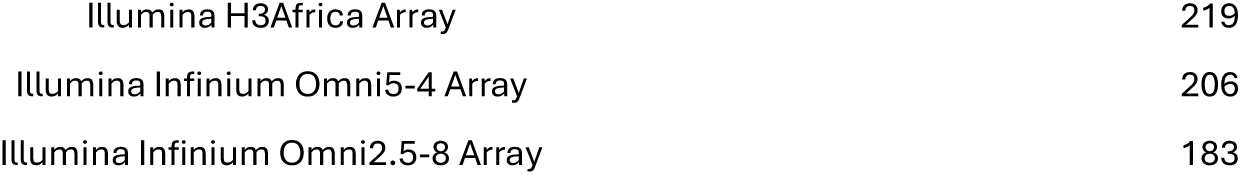
Genotype microarrays investigated in the current study.

| Genotype microarray | Number of mitochondrial markers available |
| --- | --- |
| Illumina Infinium Multi-Ethnic Global-8 Array | 774 |
| Illumina Infinium Multi-Ethnic AMR/AFR-8 Array | 643 |
| Affymetrix Genome-Wide Human SNP Array 6.0 | 410 |
| Axiom Genome-Wide PanAFR Genotyping Bundle | 261 |
| Axiom Genome-Wide Human Origins Array | 259 |
| Illumina H3Africa Array | 219 |
| Illumina Infinium Omni5-4 Array | 206 |
| Illumina Infinium Omni2.5-8 Array | 183 |

QC was performed on the genotype data using PLINK v2^27^. Individuals missing more than 10% genotypes were removed and SNPs missing more than 5% genotypes were excluded. Monomorphic SNPs were removed. SNP positions were checked to ensure genomic coordinates were mapped to the revised Cambridge Reference Sequence (rCRS)^28^. The quality controlled, unimputed files were converted to variant call format (VCF) and submitted to Haplogrep v3 for haplogroup classification^29^. Imputation was performed on the files using the MitoImpute imputation pipeline and recommended parameters (k_hap_ = 500 and MAF > 0.1%)^22^. Following imputation, monomorphic sites were excluded, and the final datasets were converted to VCF format and submitted to Haplogrep v3. The complete mitochondrial sequences were also submitted to Haplogrep v3 to obtain the true haplogroups of each sample. A flowchart outlining QC procedures and the imputation workflow is presented in Figure 2.

**Figure 2.**
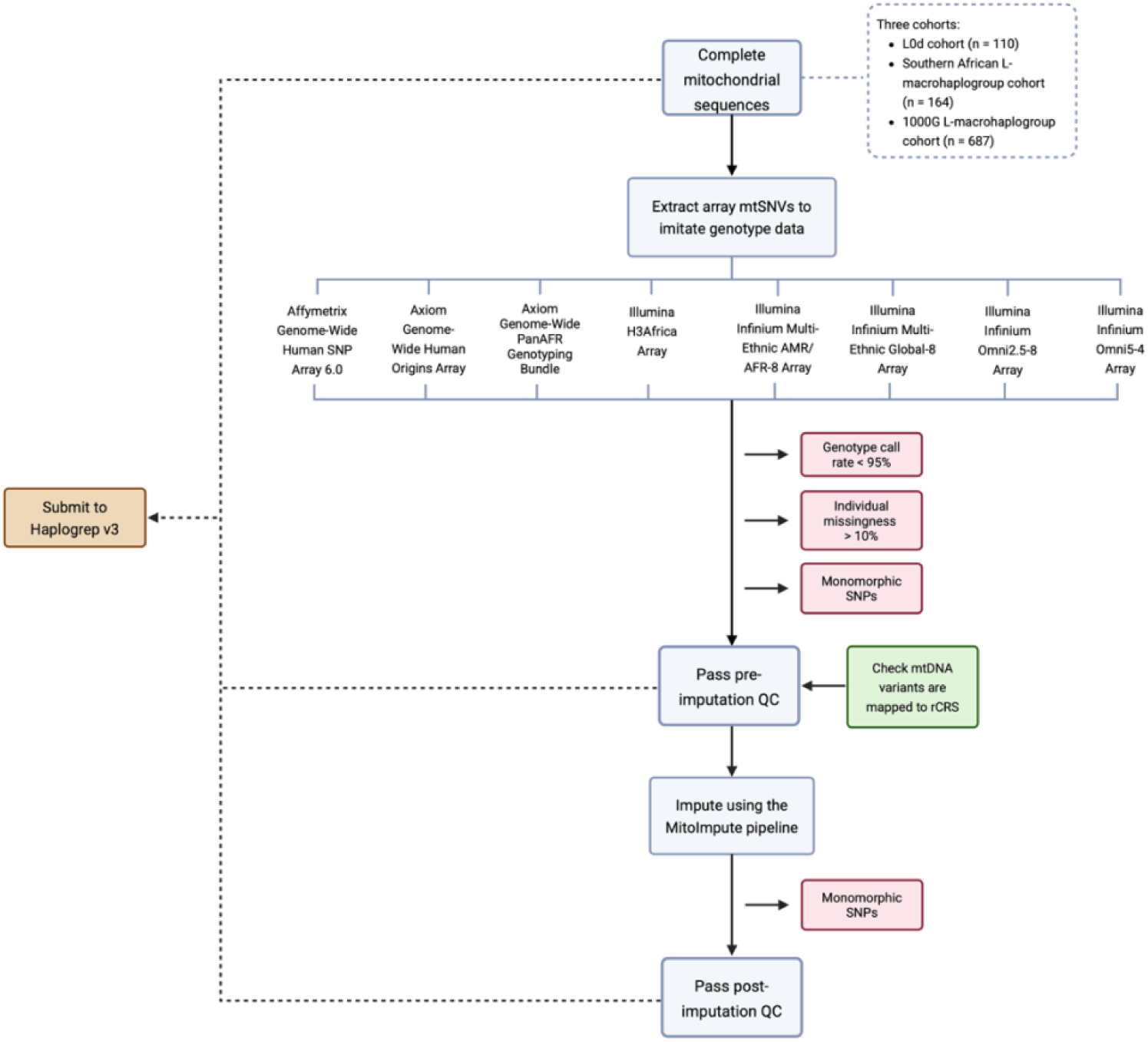
Overview of quality control (QC) and imputation workflow for assigning mitochondrial haplogroups.

Imputation performance was evaluated by assessing the similarity between imputed haplogroups and true haplogroups. For each cohort, we calculated the proportion of individuals whose assigned haplogroups matched the true haplogroups on both the array and in the imputed datasets, across four levels of haplogroup classification: 1-Level (e.g., L), 2-Level (e.g., L0), 3-Level (e.g., L0d), and 4-Level (e.g., L0d1). Wilcoxon signed-rank tests were used to determine whether imputation significantly increased the percentage of matching haplogroups at each haplogroup level (*p*-value < 0.05). Additionally, Cohen’s kappa coefficient (κ) was used to measure the level of agreement (concordance) between the haplogroups derived from array/imputed data and the true haplogroups. This coefficient accounts for agreement occurring by chance and is therefore more robust than a simple percentage agreement. Cohen’s kappa coefficient was calculated using the following formula (as implemented in the *irr* R package):

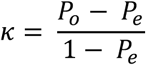

Where P_o_ is the observed agreement (i.e. the proportion of samples where the array- or imputed-derived haplogroup matches the true haplogroup) and P_e_ is the expected agreement by chance, based on the marginal probabilities of each haplogroup classification. The resulting κ value ranges from zero to one, where one indicates perfect agreement and zero indicates an agreement no better than chance. Cohen’s kappa coefficients were only determined for 4-Level haplogroups. Wilcoxon signed-rank tests were used to determine whether imputation significantly increased the level of agreement compared to array-derived haplogroups (*p*-value < 0.05).

Imputation performance was also assessed by evaluating the INFO score quality metric derived from the IMPUTE2 imputation algorithm^22,23^. Mean INFO scores and the number of imputed SNPs were calculated for each imputed dataset across eight minor allele frequency (MAF) bins (0 – 0.005, 0.005 – 0.01, 0.01 – 0.05, 0.05 – 0.1, 0.1 – 0.2, 0.2 – 0.3, 0.3 – 0.4, 0.4 – 0.5) to get a broad overview of the imputation quality across the mitochondrial genome. Additionally, since the choice of an optimal INFO score threshold value is an ongoing issue that researchers grapple with, the genotype concordance between imputed and complete mitochondrial sequence data was evaluated using BCFTools^30^ across eight different INFO score thresholds (INFO ≥ 0, INFO ≥ 0.3, INFO ≥ 0.4, INFO ≥ 0.5, INFO ≥ 0.6, INFO ≥ 0.7, INFO ≥ 0.8 and INFO ≥ 0.9). Based on these results, we aimed to provide recommendations for appropriate INFO score cutoff values for mitochondrial data. Finally, we evaluated the number of shared sites between the complete mitochondrial sequence data and the imputed data to assess the level of noise introduced by imputation, represented by the number of imputed SNPs absent in the complete mitochondrial sequence data. We also examined the extent of unique genetic diversity in the study cohorts, represented by the number of SNPs found only in the sequence data and not captured by the imputation panel. Collectively, these metrics offered a comprehensive view of the MitoImpute reference panel’s performance in our cohorts and helped determine its suitability for use in southern African population groups.

### 2.2. Harmonisation of mtDNA variants across in-house genetic datasets

Once we determined that the MitoImpute reference panel and imputation pipeline could be reliably used to infer untyped genotypes in southern African populations, we applied the pipeline to merge mitochondrial genotype datasets from southern African TB case-control cohorts and conduct an miWAS for TB susceptibility. An overview of the workflow for genotype QC, imputation and merging is provided in Figure 3. We included three microarray datasets in this analysis: cohort 1 was genotyped using the Illumina Multi-Ethnic Array and cohorts 2 and 3 were genotyped using the Illumina H3Africa Array. Cohorts 1 and 2 comprise individuals who self-identify as SAC and were recruited from the Cape Town Metropolitan area, Western Cape. Cohort 3 included self-identified SAC individuals from Upington, Northern Cape. We also included two cohorts with complete mitochondrial sequences. Cohort 4 includes 164 individuals who self-identify as isiXhosa who were recruited from the Cape Town Metropolitan area, Western Cape, and cohort 5 includes 180 individuals who self-identify as SAC who were recruited in Upington, Northern Cape. The complete mitochondrial sequences for cohort 4 and 5 were extracted from WGS using the MitoHPC pipeline^31^ (methods outlined in Supplementary Methods).

**Figure 3.**
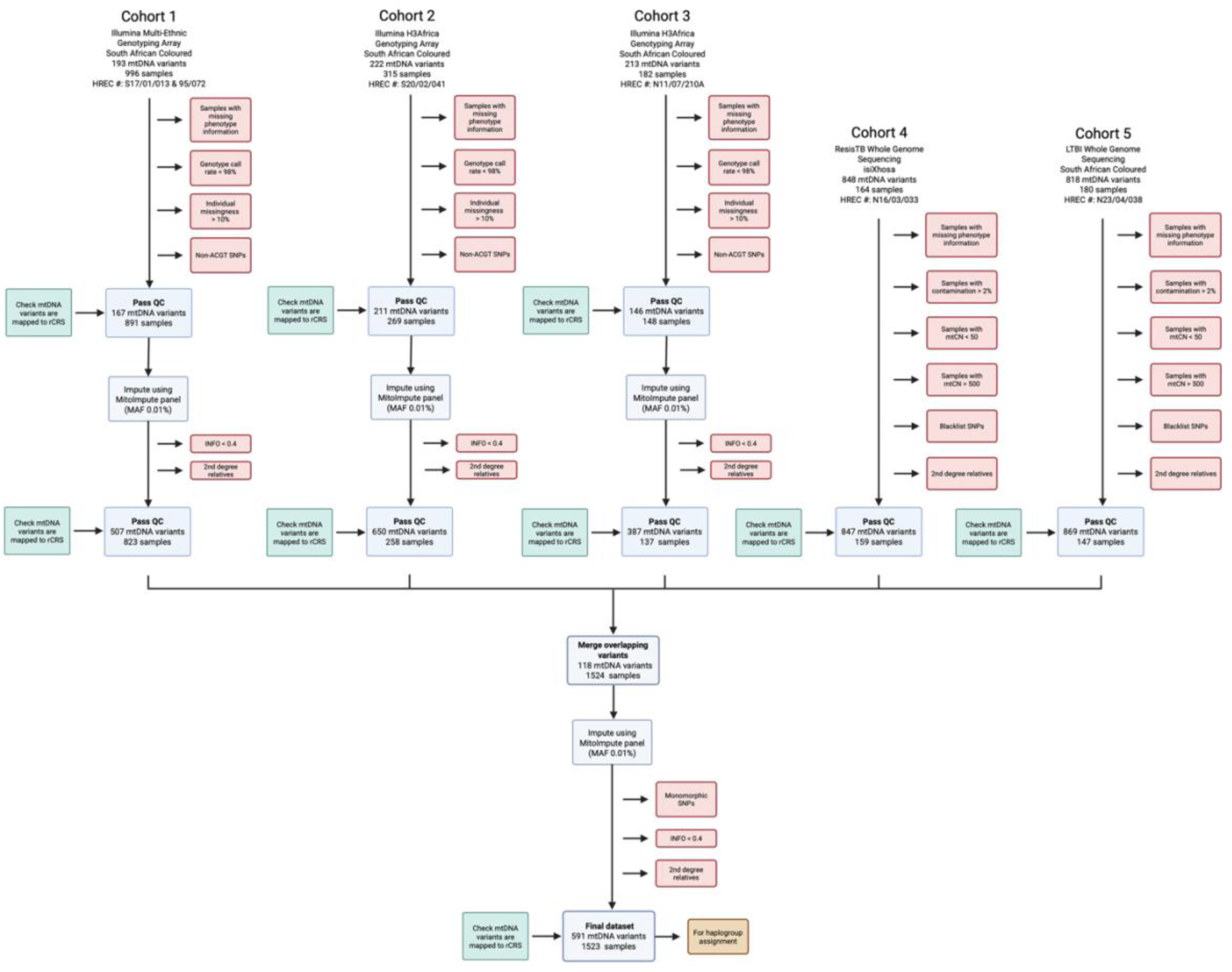
An overview of the workflow used for quality control (QC), imputation, and merging of in-house mtDNA genotype datasets.

The microarray datasets were quality controlled using PLINK v2. Samples with missing phenotype, age or sex information were removed. Samples with genotype call rates less than 90% and sites with genotype call rates less than 98% were excluded. After QC, the genotype datasets were imputed separately using the MitoImpute imputation pipeline and recommended parameters (k_hap_ = 500 and MAF > 0.1%). Following imputation, imputed variants with INFO score values less than 0.4 and monomorphic SNPs were excluded. Since there are no standard procedures for adjusting for close maternal relatedness using mitochondrial data, related individuals were identified using KING and the corresponding autosomal genotype data for each cohort^32^. Individuals identified as second-degree relatives or closer based on autosomal data were removed from the mitochondrial genotype dataset using PLINK v2.

The complete mitochondrial sequence cohorts underwent QC steps similar to those recommended by Laricchia et al. (2022)^3^. Samples with contamination levels greater than 2% or haplogroup quality scores less than 80% were excluded from further analysis. Samples with mtDNA copy number (CN) less than 50 and greater than 500 were also excluded as per previous recommendations^3^. A predefined list of artifact prone-sites (positions 301, 302, 310, 316, 3107 and 16182) were excluded to remove potential sequencing artifacts or technical errors. Individuals identified as second-degree relatives or closer based on autosomal WGS data were excluded from the mitochondrial sequence datasets.

The five cohorts were compared to identify SNPs shared across all datasets, and only these overlapping variants were merged. The merged dataset was then imputed using the MitoImpute imputation pipeline. Following imputation, we excluded imputed variants with INFO scores less than 0.4 and monomorphic variants. The final harmonised dataset contained 1523 samples with 591 mtDNA variants. Haplogroups were assigned to individuals in the final merged dataset using Haplogroup v3. Haplogroups were consolidated into broader categories (L0a, L0b, L0d, L0f, L0g, L0k, L1, L2, L3, L4, L5, M, N, and R) to enhance the stability of downstream regression models. Additionally, samples belonging to unique haplogroups (i.e., haplogroups represented by only one individual in the dataset) were excluded from further analyses, leaving a total of 1507 samples (588 TB cases and 919 controls) available for the miWAS.

### 2.3. Statistical analysis

Principal component analysis (PCA) was performed on mitochondrial SNPs in the final harmonized dataset using the *vcfR* package in RStudio (version 2025.05.0+496)^33^. Prior work established that mitochondrial PCA recapitulates haplogroup information^20^. To evaluate whether this relationship persists in our multi-ancestry southern African cohort, we estimated the proportion of variance in each principal component (PC) attributable to haplogroup using the following linear regression models (implemented in R):

1. <u>Haplogroup-only model</u> – proportion of variance explained by haplogroup alone:

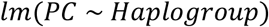
2. <u>Cohort-only model</u> – the proportion of variance explained by cohort alone:

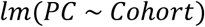
3. <u>Haplogroup-cohort model</u> – the proportion of variance explained by both cohort and haplogroup:

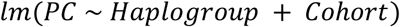

Results from these linear regression tests were used to determine whether mitochondrial PCs successfully reflect the population structure reflected in haplogroup assignment in a complex cohort.

Finally, we performed a logistic regression test to identify mtDNA variants associated with TB status. Previous work has shown that incorporating autosomal PCs provides little additional control for population structure or confounding beyond what is achieved using mitochondrial PCs^20^. Therefore, we did not determine nuclear PCs for our cohort. However, we wanted to confirm that mitochondrial PCs provide better control for population structure than haplogroups^20^. To do so, we ran two miWAS analyses. In the first miWAS, sex, age, cohort and haplogroup were included as covariates, and the logistic regression model was implemented in R according to the following formula:

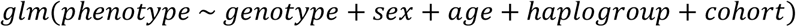

In the second miWAS, sex, age, cohort and the first 5 mitochondrial PCs were included as covariates:

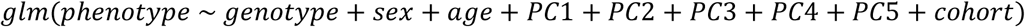

The genomic inflation coefficients (λ) were compared for both models to determine which strategy provided better control for population stratification. For both models, we used a Bonferroni-adjusted significance threshold to correct for multiple testing^34^. First, we performed a spectral decomposition analysis, using the method developed by Galwey^35^, to determine the effective number of independent genetic tests (M_eff_). Since the mitochondrial genome is inherited as a discrete unit without recombination, mtDNA variants are highly correlated. Therefore, estimating M_eff_ for Bonferroni allows for a more realistic multiple-testing correction than using the full number of variants tested, which would be overly conservative. Briefly, M_eff_ was calculated from the eigenvalues of the variant correlation matrix. M_eff_ was then divided by the threshold for significance (alpha, α = 0.05) to obtain the Bonferroni-corrected threshold. The M_eff_ calculated from the final merged dataset (which contained 591 mtSNVs) was 109. Hence, the Bonferroni-corrected significance threshold (α_Bonferroni_) was 0.00046 (rounded up to 0.0005).

### 2.4. Ethics statement

This study was approved by the Health Research Ethics Committee (HREC) of Stellenbosch University (protocol number S24/03/058). Cohorts 1–5 comprised pre-existing genetic datasets, each of which had received independent ethical approval as part of their respective studies (HREC approval numbers provided in Figure 3). All participants had provided informed consent for the use of their genetic data in future research on TB host genetics.

## 3. Results

### 3.1. Imputation performance

To assess the suitability of the MitoImpute reference panel for southern African cohorts, we first determined whether imputed haplogroups more closely aligned with the true haplogroups (derived from complete mitochondrial sequences) compared to those assigned using array data. The proportion of individuals whose array- and imputed-derived haplogroups matched the true haplogroups was calculated for all three cohorts (Supplementary Data: Table 2a-c) with results shown in Figure 4. Wilcoxon signed-rank tests for significance were calculated at each haplogroup level for each cohort (Supplementary Data: Table 3). While imputation did not improve haplogroup classification at lower haplogroup levels (1-Level and 2-Level), it did improve classification at higher levels (3-Level and 4-Level; *p*-values ranged from 0.0156 to 0.036). Thus, imputed mitochondrial data enabled more accurate classification for higher-level haplogroups compared to array-based data. The quality of haplogroup assignment also depends on the number of haplogroup-defining markers genotyped prior to imputation. The Illumina Infinium Multi-Ethnic Global-8, the Illumina Infinium Multi-Ethnic AMR/AFR-8, and the Affymetrix Genome-Wide Human SNP Array 6.0 include a greater number of L-macrohaplogroup defining SNPs compared to the other arrays assessed (Supplementary Data: Table 4a and 4b). Consequently, these arrays yielded better quality haplogroup assignments both before and after imputation.

**Figure 4.**
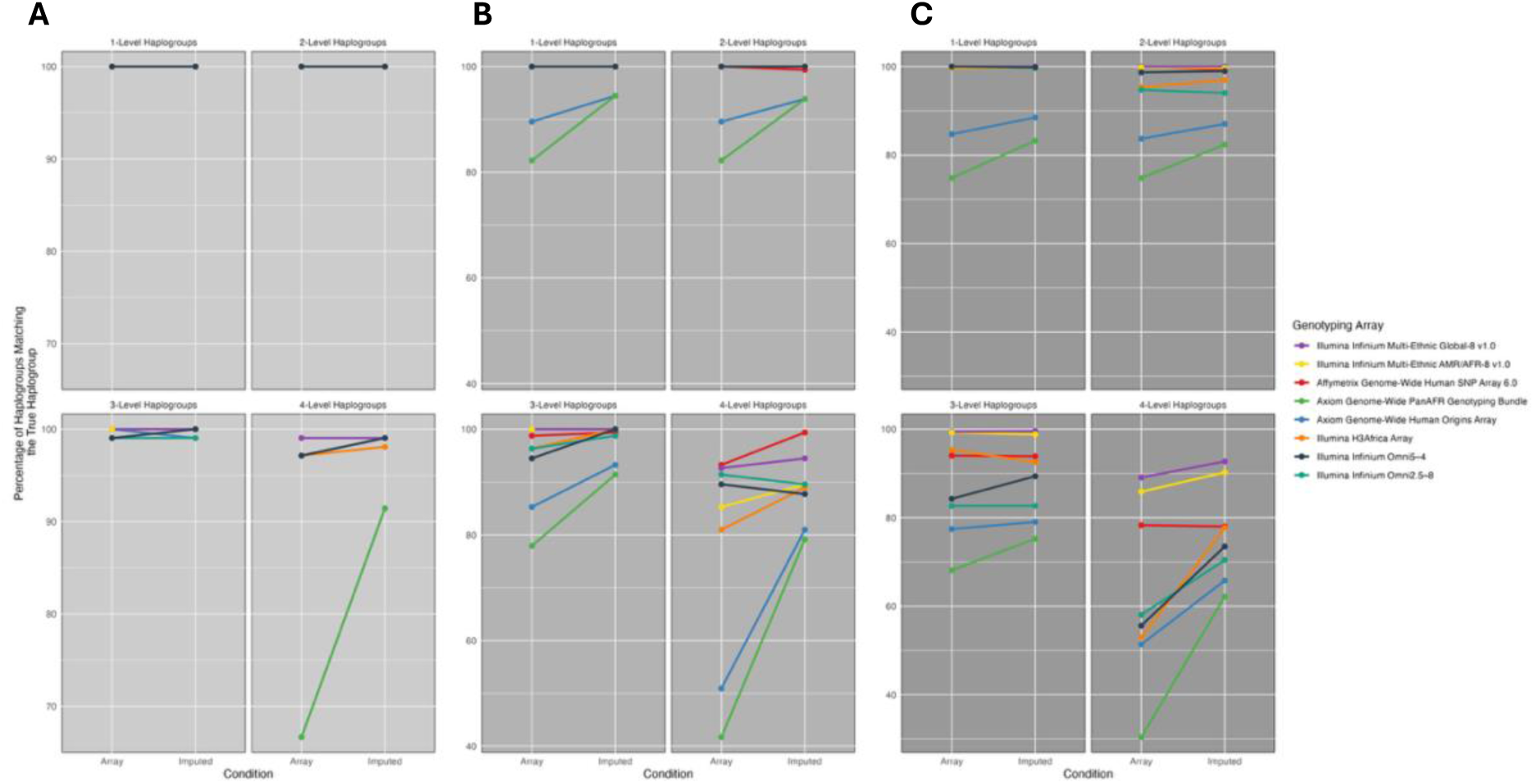
Percentage of array- and imputed-derived haplogroups that match the true haplogroup across four levels of haplogroup classification for: **(A)** the southern African L0d cohort (n = 110), **(B)** the southern African L-macrohaplogroup cohort (n = 164), and **(C)** the 1000G Phase 3 L-macrohaplogroup cohort (n = 687). The figure legend is ordered by the number of markers available on each genotype microarray before imputation, with the Illumina Infinium Multi-Ethnic Global-8 v1.0 array having the most and the Illumina Infinium Omni2.5-8 array the fewest.

Cohen’s kappa coefficients were used to assess the level of agreement between true haplogroups and array- or imputed-derived haplogroups. Coefficients were calculated only for 4-Level haplogroups across all three cohorts (Supplementary Data: Table 5). Generally, haplogroups derived from imputed data showed larger kappa coefficients than those derived from array data (Figure 5). This improvement was statistically significant in the southern African L0d cohort (*p-*value = 0.0213) and the 1000G Phase 3 L-macrohaplogroup cohort (*p-*value = 0.01563), but not in the southern African L-macrohaplogroup cohort (*p-*value = 0.07813) (Supplementary Data: Table 6). Nevertheless, within the southern African L-macrohaplogroup cohort, six of the eight genotype array platforms still showed increased kappa coefficients for imputed-derived haplogroups. Together, these findings suggest that, for higher-level haplogroup classifications, imputation can provide more accurate haplogroup assignments than array-based data.

**Figure 5.**
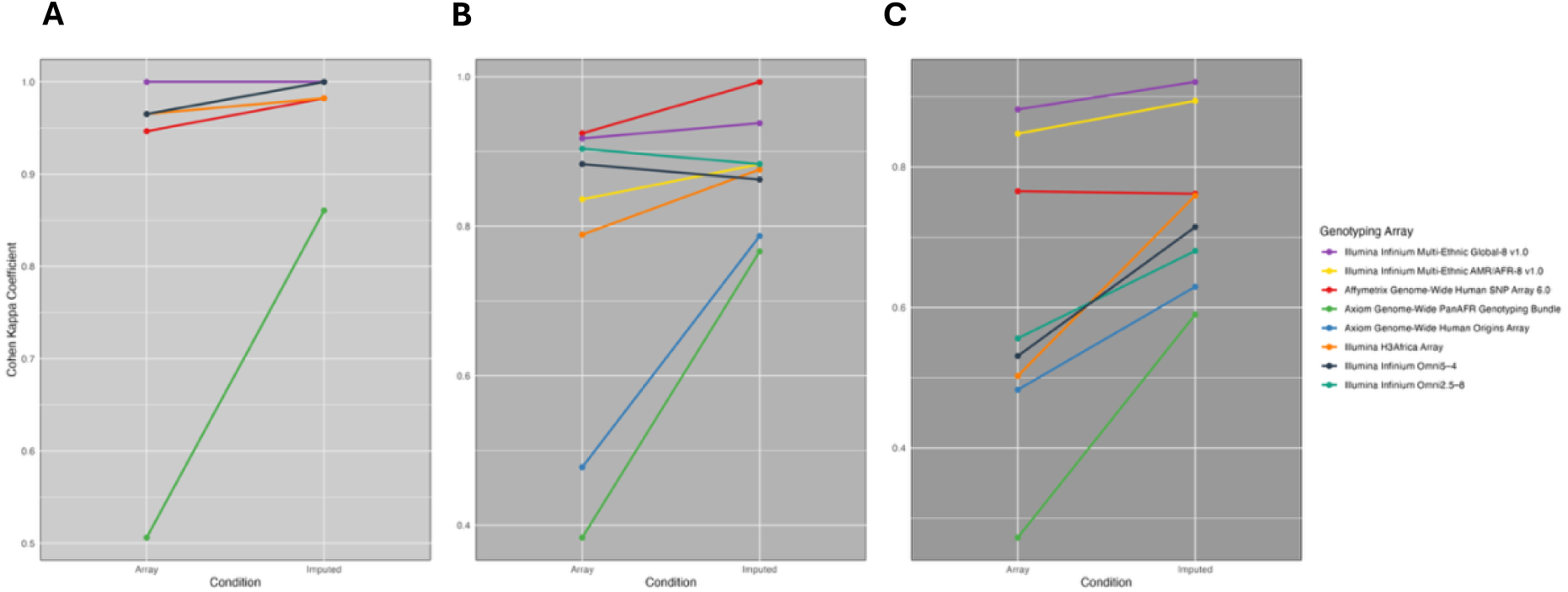
Differences in the Cohen’s kappa coefficients (i.e., the level of agreement) between array- or imputed-derived haplogroups and the true haplogroups (for 4-Level haplogroups only) for: **(A)** the southern African L0d cohort (n = 110), **(B)** the southern African L-macrohaplogroup cohort (n = 164), and **(C)** the 1000G Phase 3 L-macrohaplogroup cohort (n = 687). The figure legend is ordered by the number of markers available on each genotype microarray before imputation, with the Illumina Infinium Multi-Ethnic Global-8 v1.0 array having the most and the Illumina Infinium Omni2.5-8 array the fewest.

Additionally, we assessed the quality and number of imputed SNPs in each cohort (Supplementary Figures 3 and 4). As with autosomal data, imputation quality for mtDNA variants depends on the number of markers initially genotyped. Across all three cohorts assessed, the microarrays with the highest marker density (Illumina Infinium Multi-Ethnic Global-8 v1.0, the Illumina Infinium Multi-Ethnic AMR/AFR-8 v1.0 and the Affymetrix Genome-Wide Human SNP Array 6.0) achieved higher mean INFO scores across all MAF bins compared to the other arrays evaluated (Supplementary Figure 3). Imputation quality was lowest for rare variants (MAF < 0.01), consistent with patterns observed in autosomal data. However, imputation quality did not consistently increase with MAF. Notably, a marked drop in mean INFO score was observed for the 1000G Phase 3 L-macrohaplogroup cohort at relatively high MAFs (Supplementary Figure 3C). A smaller drop in mean INFO score was also observed in the southern African L-macrohaplogroup cohort at the 0.2-0.3 MAF bin (Supplementary Figure 3B). In both cases, the sharp decrease in mean INFO score is likely due to fewer markers being imputed in this MAF range overall (Supplementary Data: Table 7b-c). Most imputed SNPs fall within the lowest MAF bin (0 – 0.005) (Supplementary Figure 4; Supplementary Data: Table 7a-c), a distribution pattern that is consistent with observations from autosomal data^36^.

Selecting an appropriate INFO score cutoff remains a challenge for researchers. To determine the impact of different cutoff thresholds on concordance with true genotypes, we examined the genotype concordance across increasing INFO score cutoffs (Figure 6). Additionally, we examined the number of SNPs retained in the dataset at each cutoff (Figure 7). Given that genotype concordance plateaued at approximately 90% for variants with INFO ≥ 0.4, this threshold was selected for downstream analyses. This cut-off provides an optimal trade-off between retaining a substantial number of variants while filtering out low-quality variants that could introduce noise or spurious associations.

**Figure 6.**
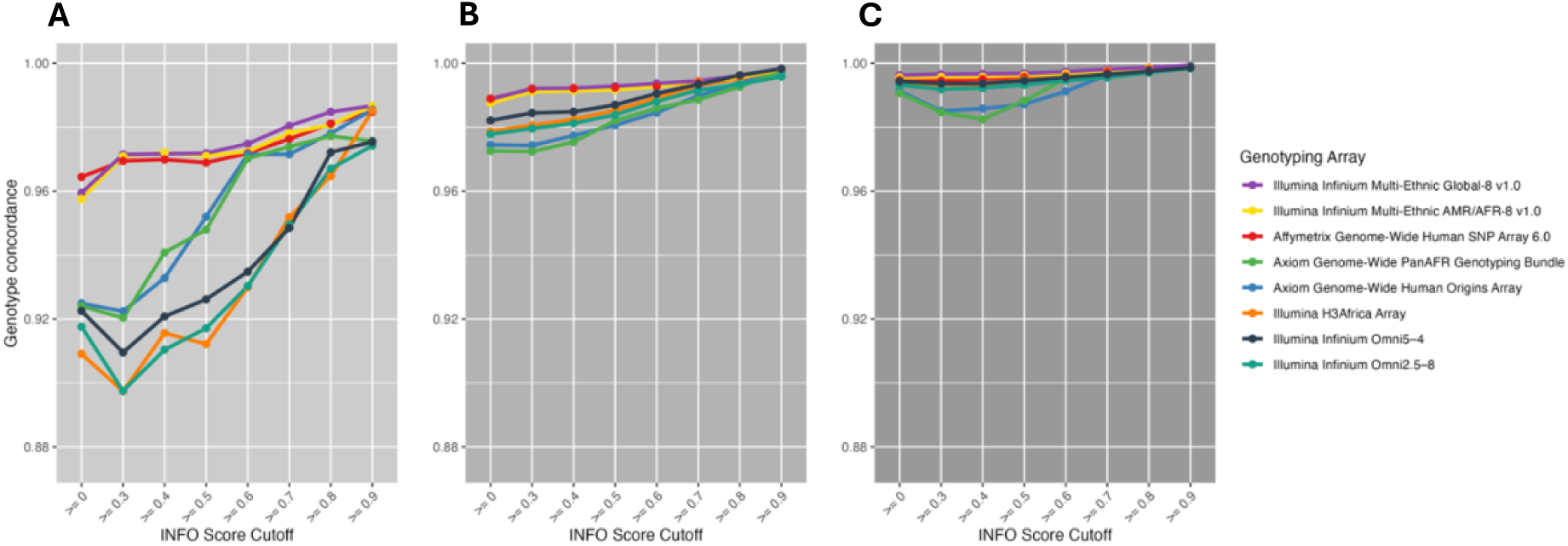
Genotype concordance between imputed and sequenced datasets across all INFO score cutoff thresholds for: **(A)** the southern African L0d cohort (n = 110), **(B)** the southern African L-macrohaplogroup cohort (n = 164), and **(C)** the 1000G Phase 3 L-macrohaplogroup cohort (n = 687). The figure legend is ordered by the number of markers available on each genotype microarray before imputation, with the Illumina Infinium Multi-Ethnic Global-8 v1.0 array having the most and the Illumina Infinium Omni2.5-8 array the fewest.

**Figure 7.**
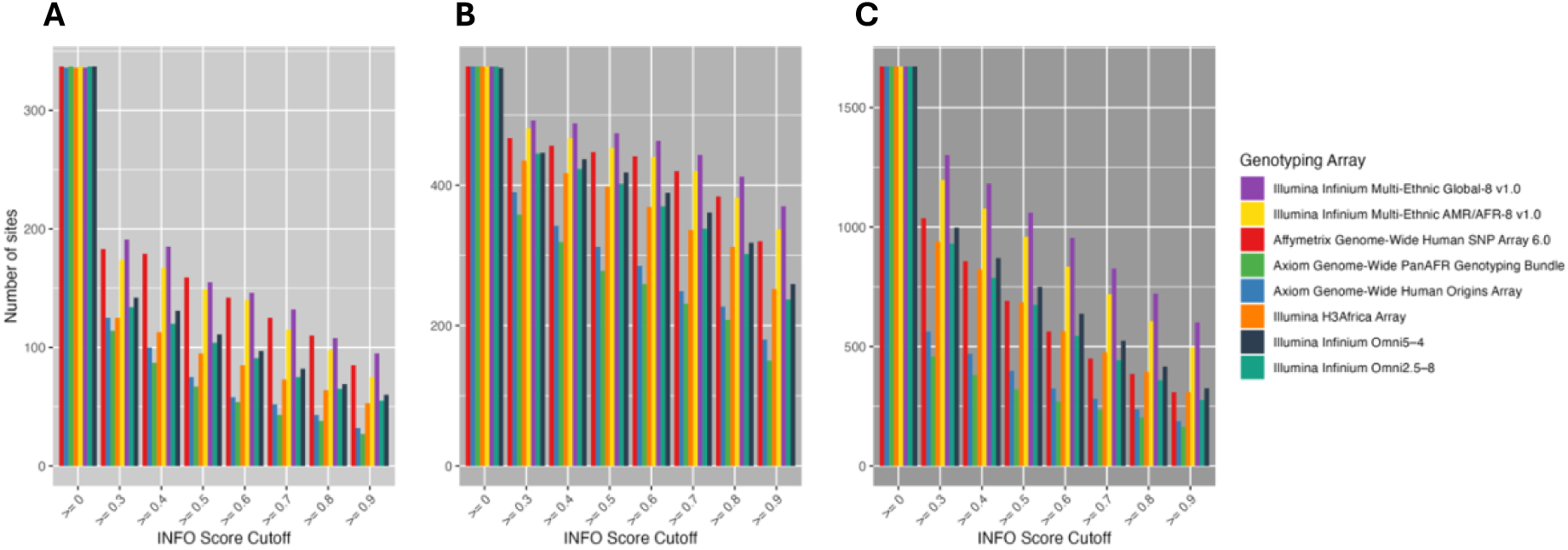
Distribution of the number of imputed SNPs across all INFO score cutoff thresholds for **(A)** the southern African L0d cohort (n = 110), **(B)** the southern African L-macrohaplogroup cohort (n = 164), and **(C)** the 1000G Phase 3 L-macrohaplogroup cohort (n = 687). The figure legend is ordered by the number of markers available on each genotype microarray before imputation, with the Illumina Infinium Multi-Ethnic Global-8 v1.0 array having the most and the Illumina Infinium Omni2.5-8 array the fewest.

The number of shared sites between the complete mitochondrial sequences and the imputed data are shown for each cohort is depicted in Supplementary Figures 5-7 (see also Supplementary Data: Table 9). Overall, the proportion of SNPs that were unique to the imputed data is relatively low, ranging from 8.31 – 17.24% of imputed sites. In contrast, the number of sites unique to the sequenced data is substantially higher, highlighting the remarkable genetic diversity that can be captured by complete mitochondrial genome sequencing. While the MitoImpute reference panel includes over 5 000 sequences from macrohaplogroups L0 – L5 (representing more than 13% of the entire panel), the unique genetic diversity in mitochondrial sequences from African populations mean that many variants present in these sequences are not represented in the reference panel and thus cannot be imputed. Consequently, complete mtDNA sequencing remains the most reliable approach for rare and/or novel variant discovery in African populations. However, following our evaluation of the performance of the MitoImpute reference panel, we determined that this imputation reference panel is suitable for the imputation of common and haplogroup-defining mtDNA variants in southern African populations.

### 3.2. Summary statistics of the harmonized mitochondrial dataset

The final dataset generated from the steps outlined in Figure 5 included 1507 samples (588 TB cases and 919 controls) with 591 mtDNA SNPs. Female samples were disproportionately over-represented in the dataset, particularly in the case group (χ^2^ test *p*-value = 2.7×10^-8^) (Supplementary Figure 8A). On average, cases were older than controls (Wilcoxon test *p*-value = 1.74×10^-6^) (Supplementary Figure 8B). Haplogroup distribution between cases and controls did not differ significantly (Fisher’s Exact test *p*-value = 0.05) (Figure 8A). However, there was a significant difference in the distribution of haplogroups among the five individual cohorts (Fisher’s Exact test *p*-value = 0.0005) (Figure 8B). Moreover, the distribution of cases and controls differed significantly among the five different cohorts (Supplementary Figure 9). The substantial variation in phenotype and haplogroup distributions across cohorts required including “cohort” as a covariate in the miWAS to account for potential batch effects and minimize spurious associations.

**Figure 8.**
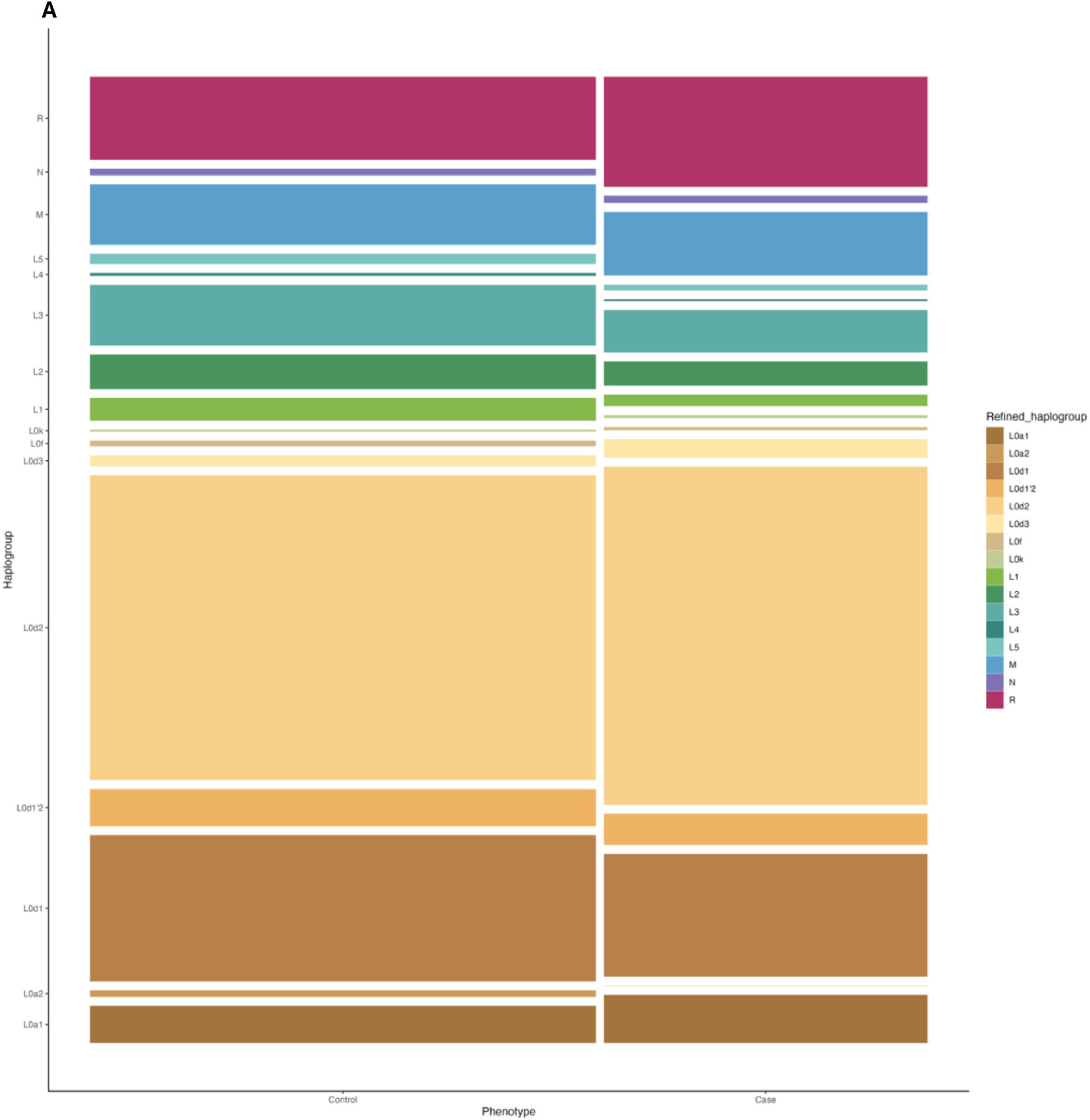
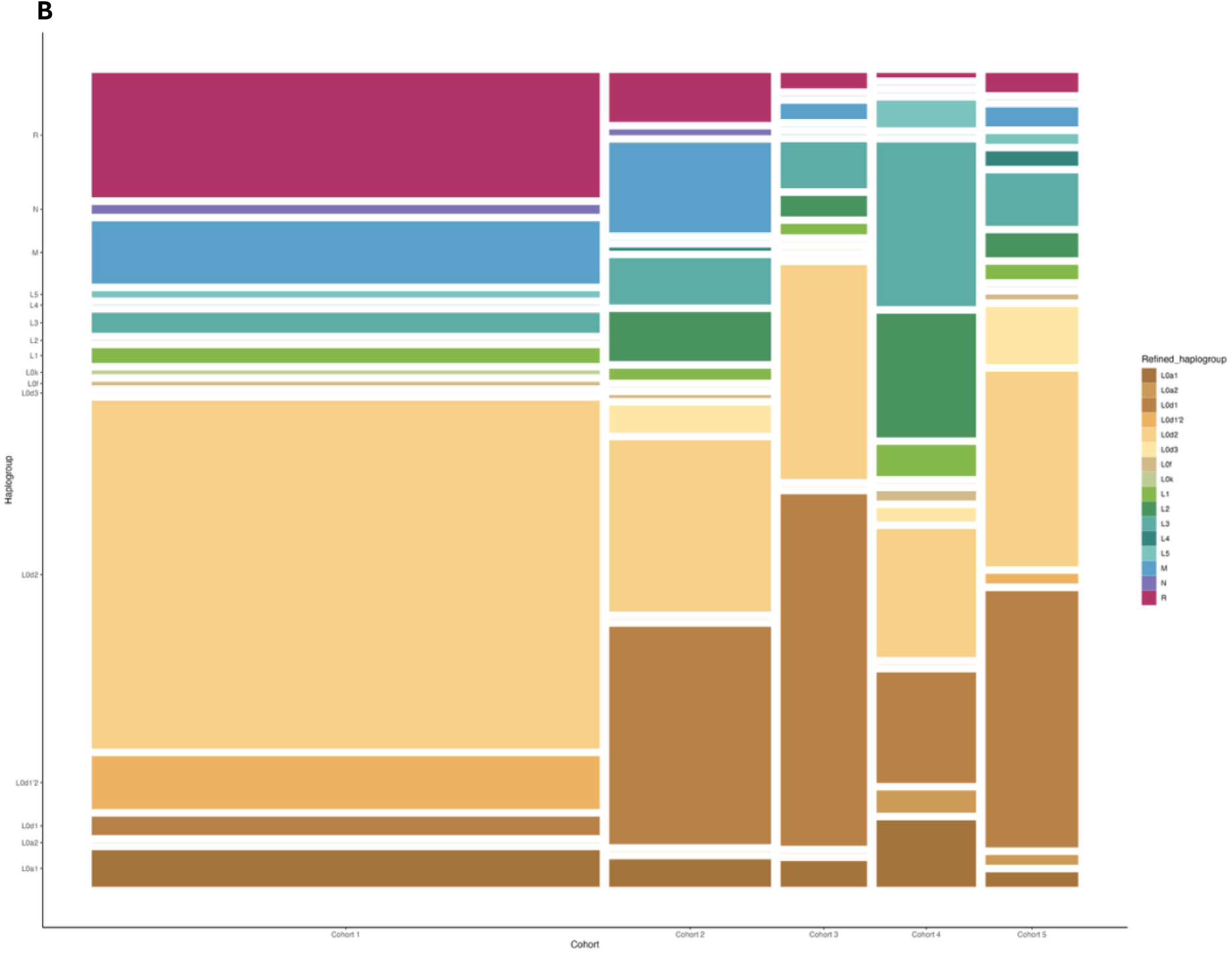
**(A)** Distribution of haplogroups between cases and controls in the merged dataset. The figure legend lists the haplogroups represented in the dataset, ordered alphabetically and numerically. The difference in haplogroup distribution between cases and controls is not statistically significant, as determined by Fisher’s Exact test (*p*-value > 0.05). **(B)** Distribution of haplogroups among the five different cohorts that make up the merged dataset. The figure legend lists the haplogroups represented in the dataset, ordered alphabetically and numerically. The difference in haplogroup distribution among the cohorts is statistically significant, as determined by Fisher’s Exact test (*p*-value < 0.05). Cohort 1 and 2 comprise self-identified SAC individuals recruited from the Cape Town Metropolitan area in the Western Cape. Cohort 3 and 5 comprise self-identified SAC individuals recruited from Upington and surrounding areas in the Northern Cape. Cohort 4 comprises self-identified Xhosa individuals recruited from the Cape Town Metropolitan area in the Western Cape.

The PCA plot for mtDNA variants (Figure 9A) demonstrates clustering of samples by ancestry, which closely parallels the patterns of haplogroup divergence observed in the mitochondrial phylogenetic tree. It illustrates the extensive genetic diversity observed among the early diverging haplogroups (L0d, L0f, L0k and L0a), as well as the distinct genetic bottleneck following the migration of anatomically modern humans out of Africa, reflected by the clustering of L3 and non-African haplogroups (M, N and R). Within our harmonized dataset, the L0d2 haplogroup is present at the highest frequency (∼38%), followed by L0d1 (∼17%) (Figure 9B). Additionally, the cohort (consisting primarily of self-identified SAC individuals) exhibits remarkable haplogroup diversity, encompassing nearly all major mitochondrial lineages. This diversity reflects the complex admixture and demographic history of this population. When samples are color-coded by cohort, they are distributed relatively even across the PCs, with no evidence of cohort-driven clustering (which would suggest the presence of severe batch effects) (Supplementary Figure 10).

**Figure 9.**
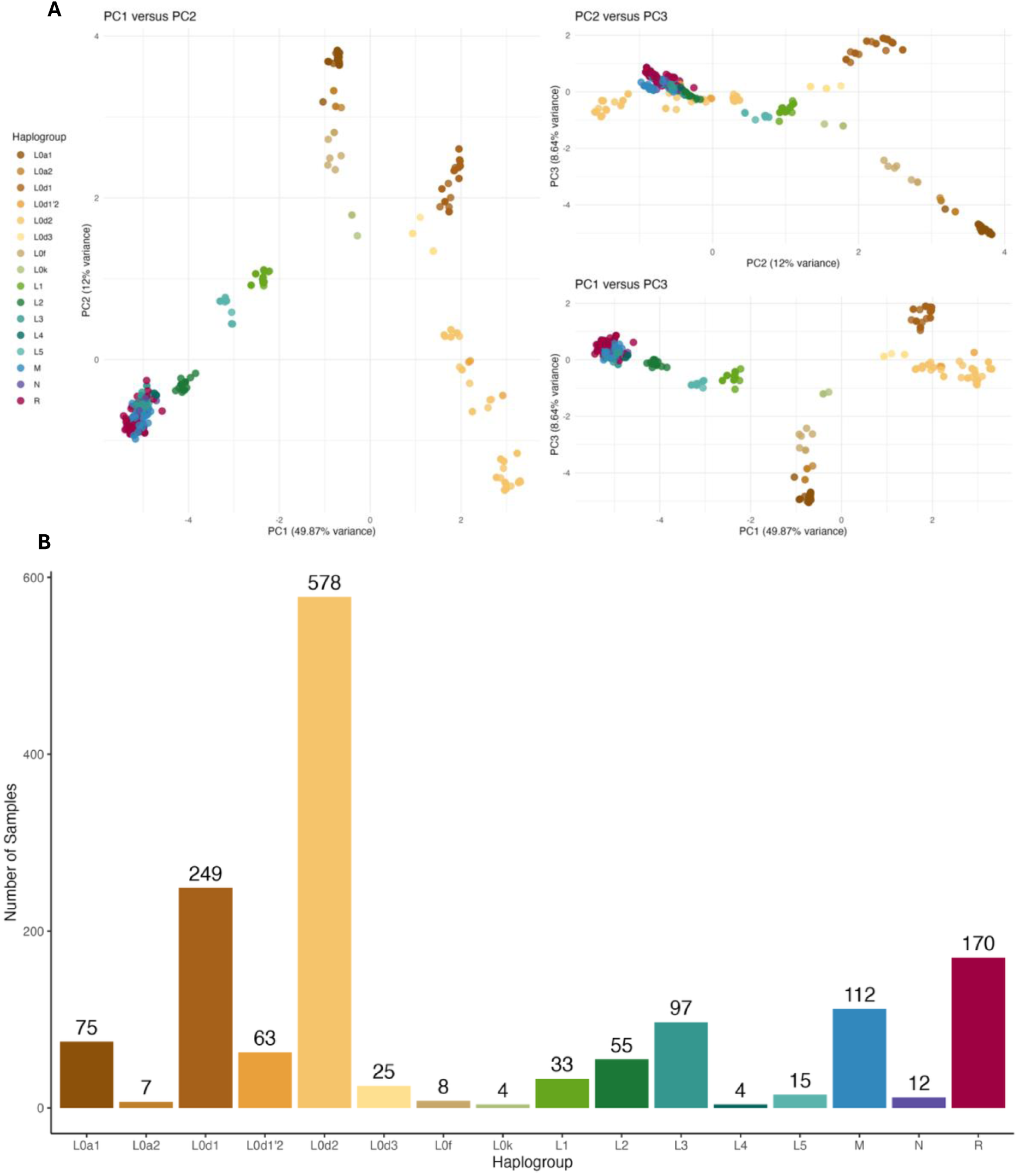
**(A)** PCA for mtDNA variants with PC1, PC2 and PC3 plotted against each other. Samples are color-coded by haplogroups. **(B)** Distribution of haplogroups in the merged dataset.

To determine whether mitochondrial PCs reflect haplogroup information, linear regression models were used to estimate the proportion of variance in each of the first five PCs attributable to haplogroups. Haplogroups accounted for over 90% of the variance across all PCs 1-5 (Table 2), suggesting that mitochondrial PCs robustly capture haplogroup structure.

**Table 2.** Results for the linear regression tests estimating the proportion of variance in PCs 1-5 explained by haplogroup, cohort or both. The proportion of variance is represented by the coefficient of determination, R^2^. Residual variance (i.e. variance that is not explained by either haplogroup or cohort) is also shown.

| Principal component | $R^2$ by haplogroup (%) | $R^2$ by cohort (%) | $R^2$ by haplogroup + cohort (%) | Residual variance (%) |
| --- | --- | --- | --- | --- |
| PC1 | 94.5 | 3.3 | 94.6 | 5.4 |
| PC2 | 97.8 | 9.6 | 97.8 | 2.2 |
| PC3 | 97.5 | 3.9 | 97.9 | 2.1 |
| PC4 | 96.4 | 7.3 | 96.4 | 3.6 |
| PC5 | 93.5 | 4.5 | 93.6 | 6.4 |

### 3.3. Mitochondrial genome-wide association results

To assess whether mitochondrial PCs offered improved correction for population structure relative to haplogroup-based adjustment, we compared the genomic inflation coefficients (λ) from two miWAS models. Both models showed mild deflation, with λ = 0.842 for the PC-adjusted analysis and λ = 0.844 for the haplogroup-adjusted analysis, indicating comparable control of test statistic inflation. Consistent with these values, the corresponding QQ-plots (Supplementary Figures 12A and 12B) show slight deflation. In the miWAS adjusted for mitochondrial PCs, two variants approached (but did not surpass) the genome-wide significance threshold (Figure 10). No variants exceeded the significance threshold in the haplogroup-adjusted model. Full association results for both models are provided in Supplementary Tables 10a and 10b. Overall, these findings suggest that mitochondrial PCs and haplogroup assignments perform similarly in accounting for underlying population structure in this highly diverse cohort, although neither model revealed statistically significant associations with TB susceptibility. The power to detect significant associations depends on sample size, MAF and variant effect size, typically expressed as odds ratios (OR) for binary traits in logistic regression tests. For our miWAS of TB susceptibility, a sample size of 1507 (comprising 588 TB cases and 919 controls) has sufficient power (≥ 80%) to detect common variants (MAF ≥ 10%) with modest effect sizes (OR ≥ 1.4) (Supplementary Figure 13). However, the detection of variants with small effect sizes is limited in our study and would necessitate a larger sample size.

**Figure 10.**
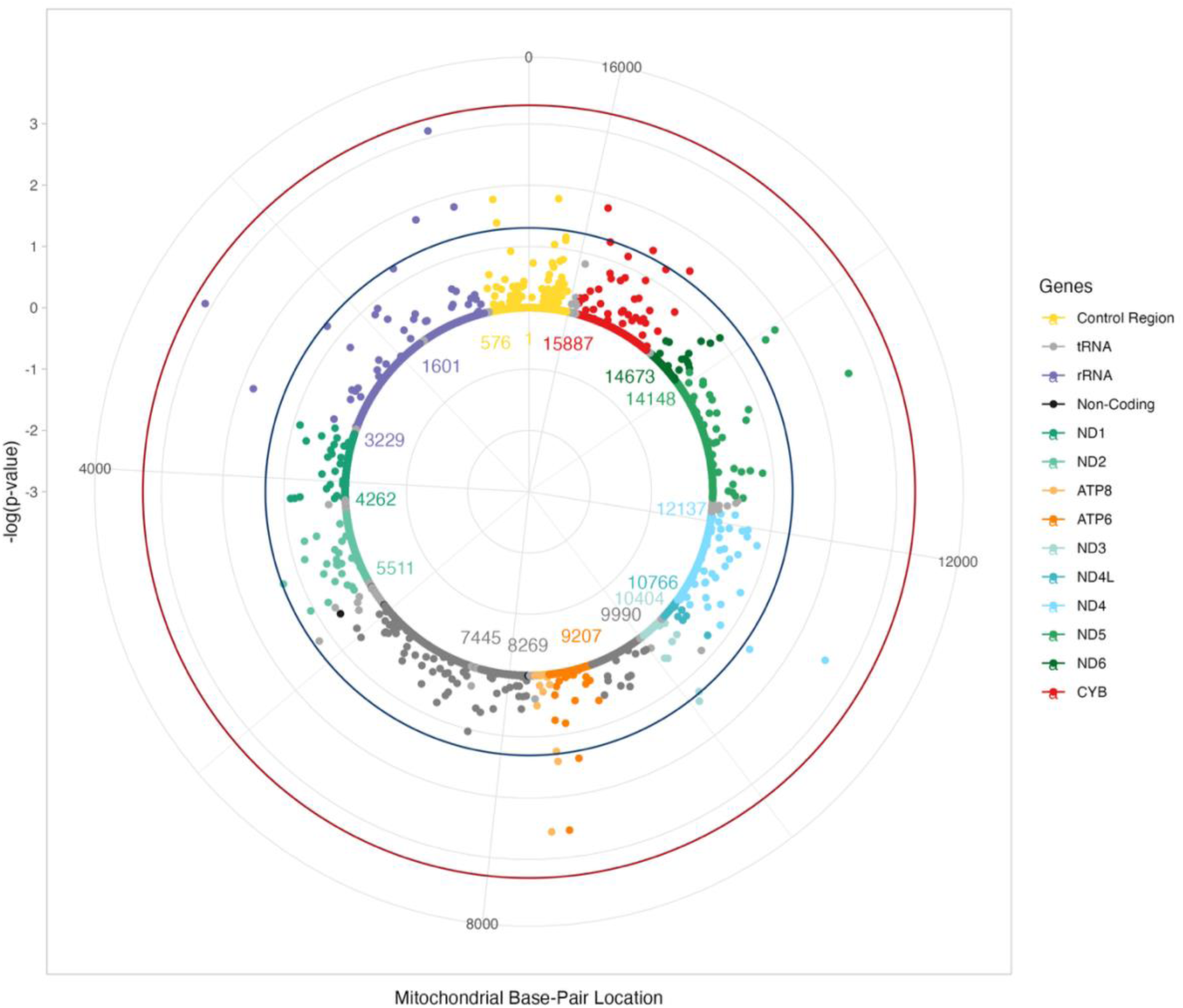
Circular Manhattan plot showing the log transformation of *p*-values obtained for the PC-adjusted miWAS model. Each dot represents a mitochondrial variant, color-coded by gene. The base pair positions are provided on a circular axis to represent the circular nature of the mitochondrial genome. The −log10(*p*-values) are provided on the radial y-axis. The inner blue circle represents the *p-*value threshold of 0.05 and the outer red circle represents the Bonferroni-adjusted *p*-value threshold of 0.0005.

## 4. Discussion and Conclusions

The persistent lack of southern African representation in genetic studies and reference databases poses a critical limitation for the field. When populations with extensive genetic diversity are excluded from the development of new tools, including imputation panels, the resulting methods may fail to perform reliably across global populations or in cohorts with complex population structure. This underrepresentation not only overlooks a substantial proportion of human genetic variation but also risks producing methods that are insufficiently generalizable. In this study, we conducted a comprehensive evaluation of the publicly available MitoImpute reference panel to determine its suitability for use in southern African populations. While the panel performed well in enabling accurate haplogroup prediction (Figures 4 and 5) and successfully imputing a proportion of mtDNA genotypes across cohorts (Figure 6), a substantial amount of genetic variation was not captured (Supplementary Figures 5–7). This uncaptured diversity likely includes population-specific mtSNVs that may contribute to complex traits. Consequently, reliance on imputed genotypes derived from a panel with limited southern African representation may constrain the ability to detect biologically meaningful associations in these populations.

Increasing the representation of southern African sequences in reference panels is particularly valuable, as global populations encompass only a subset of the genetic diversity present in sub-Saharan Africa, and such inclusion is likely to improve imputation accuracy across many different global populations. Moreover, the phylogenetic nature of mitochondrial sequences could be leveraged to enhance imputation performance^37^. The MitoImpute pipeline currently relies on an imputation algorithm designed for autosomal data, using the IMPUTE2 X-chromosome function to handle haploid mitochondrial genomes. However, the mitochondrial genome has additional unique features, such as being circular rather than linear, that are not fully accounted for by this approach. The development of a phylogenetically aware imputation algorithm capable of handling circular genomes would represent a valuable advancement in mitochondrial genetics, although such development is beyond the scope of the present study.

Although not without limitations, the MitoImpute panel was instrumental in the mtDNA variant harmonisation process in this study, enabling the successful integration of five mtDNA genotype datasets to generate a larger cohort for a miWAS of TB susceptibility. Our harmonised dataset was predominantly composed of individuals from the SAC population, which has historically received contributions from multiple ancestries, including Khoe-San, Bantu-speaking southern African, European, and East and Southeast Asian^38–42^. This complex ancestral history is reflected in the extensive mitochondrial haplogroup diversity observed in our cohort (Figure 9). Given the highly heterogeneous mitochondrial population structure in southern African cohorts, careful adjustment for population stratification is essential. In this study, we demonstrated that mitochondrial PCs effectively capture underlying haplogroup structure and provide robust correction for stratification in miWAS analyses, consistent with previous observations^20^. In our miWAS of TB susceptibility, no mtSNVs surpassed the significance threshold, and the genomic inflation factors (λ = 0.842 for the PC-adjusted model; λ = 0.844 for the haplogroup-adjusted model) and QQ-plots indicated mild deflation. However, our λ estimates closely align with the genomic inflation factor reported for the UK Biobank miWAS of tuberculosis (λ ≈ 0.83), despite their analysis including a substantially larger sample size. Collectively, our results suggest that common mitochondrial variants do not have detectable effects on TB susceptibility in these cohorts. Nevertheless, expanding sample sizes and incorporating broader African mtDNA diversity may help reveal contributions from rare or population-specific variants not captured here.

In light of these results, it is clear that larger sample sizes and fuller representation of mitochondrial variation, particularly from underrepresented African haplogroups, are critical to strengthening the power of miWAS in diverse populations. Given the limited number of population-specific mtSNVs imputed by the MitoImpute reference panel, variants unique to southern African populations that may influence TB susceptibility were likely not captured and therefore not included in this miWAS. In such cases, complete mitochondrial genome sequencing, which fully captures the breadth of genetic diversity, is recommended. Additionally, neither array-based genotyping nor imputation can recover heteroplasmic variants or copy number, which may also contribute to complex disease susceptibility; these can only be assessed using high-coverage mitochondrial sequence data.

Despite these limitations, the study has several strengths. To our knowledge, this is the first miWAS conducted for TB susceptibility and the first miWAS performed in a uniquely diverse southern African cohort. It also represents a comprehensive evaluation of a publicly available mitochondrial imputation reference panel in a highly diverse population. Moreover, we provide robust guidelines for harmonising mtDNA datasets across genotyping platforms and for appropriately accounting for population structure in a complex, multi-way ancestry mtDNA cohort. These guidelines will support future efforts to integrate larger TB case-control datasets from diverse ancestry groups to identify potential mitochondrial variants associated with TB and other complex diseases.

## 5. Acknowledgements

This study was partially funded by the South African Medical Research Council (SAMRC) Centre for Tuberculosis Research through a Competitive-collaborative Support Grant for CTR Researchers (RFA-01-052022-CSECPR). The content is solely the responsibility of the authors and does not necessarily represent the official views of the SAMRC. Dayna Croock was supported by a Stellenbosch University Postgraduate Bursary and the SAMRC Centre for Tuberculosis Research scholarship.

